# Systematic analysis of aberrances of ferroptosis reveals its potential functional roles in cancer

**DOI:** 10.1101/765826

**Authors:** Zekun Liu, Qi Zhao, Zhi-Xiang Zuo, Shu-Qiang Yuan, Kai Yu, Qingfeng Zhang, Xiaolong Zhang, Hui Sheng, Huai-Qiang Ju, Han Cheng, Feng Wang, Rui-Hua Xu, Ze-Xian Liu

**Author notes:** To whom correspondence should be addressed: Dr. Ze-Xian Liu, Dr. Rui-Hua Xu. These authors contributed equally to this work (Zekun Liu, Qi Zhao, Zhi-Xiang Zuo, Shu-Qiang Yuan).

## Abstract

Ferroptosis is a type of cell death that related to cancer, however, the characteristics of ferroptosis in cancers are still uncertain. Based on the data in The Cancer Genome Atlas, we found that most ferroptosis regulator genes (FRGs) were differentially expressed in tumors, copy number alterations (CNA) and DNA methylation contributed to their aberrant expression. We established the ferroptosis potential index (FPI) to reveal the functional roles of ferroptosis and noticed that the FPI was higher in tumors than in normal tissues in most cancers, and was associated with subtypes and clinical features. The FPI was negatively correlated with several metabolism pathways but positively associated with several important metastasis-related pathways and immune-related pathways. Higher FPI predicted worse prognosis in several tumors, while FPI and FRGs impacted drug sensitivity. Our study presents a systematical analysis of ferroptosis and its regulatory genes, and highlights the potential of ferroptosis-based cancer therapy.

## Introduction

As a newly discovered type of programmed cell death, ferroptosis results from the accumulation of iron-dependent lipid hydroperoxides and leads to cytological changes, for which the features and mechanisms are different compared to typical cell death such as apoptosis (Dixon et al., 2012). Previous studies demonstrated that during the process of ferroptosis, sufficient and approachable cellular iron was required (Dixon et al., 2012). Inhibition of system X_C_^-^, which is a membrane Na^+^-dependent cysteine-glutamate exchange transporter, and GPX4 can disrupt the oxidation-reduction balance and cause overwhelming lipid peroxidation that ultimately results in cell death (Stockwell et al., 2017; Yang et al., 2014; Yu et al., 2017). The initiation and execution of ferroptosis are affected by multiple factors including amino acids, lipids, and iron metabolism, and regulated by various signaling pathways such as amino acid and glutathione metabolism, lipid metabolism, iron metabolism, and mevalonate pathway (Stockwell et al., 2017).

As the understanding of ferroptosis has increased, its complex biological function has been revealed (Matsushita et al., 2015; Yang and Stockwell, 2016). Furthermore, ferroptosis has been found to be closely related with various human diseases including periventricular leukomalacia, Huntington’s disease and acute kidney injury (Friedmann Angeli et al., 2014; Inder et al., 2002; Linkermann et al., 2014; Skouta et al., 2014). Besides, the literatures have confirmed that ferroptosis could suppress tumor growth and kill tumor cells (Yu et al., 2017), and then plays an important role in cancers such as renal cell carcinomas and liver cancer (Sun et al., 2016b; Yang et al., 2014). For example, erastin-induced ferroptosis decreases the growth of tumors formed from human CRC cells (Xie et al., 2017). Ductal pancreatic cancer cells with a mutant *KRAS* gene were more susceptible to ferroptosis than wild type (Eling et al., 2015; Yu et al., 2017). Conversely, inducing tumor cell ferroptosis by small molecules has become an important strategy for the treatment of many tumors, such as hepatocellular carcinoma, kidney cancer, and pancreatic cancer (Eling et al., 2015; Louandre et al., 2013; Louandre et al., 2015; Yang et al., 2014). In addition, changes in the gene expression of tumor cells also affect ferroptosis, and a number of genes have been confirmed to regulate ferroptosis. For example, *ACSL4* participates in the biosynthesis and remodeling of polyunsaturated fatty acid-Pes, and the downregulation of *ACSL4* increases the resistance to ferroptosis (Dixon et al., 2015; Doll et al., 2017). Furthermore, ferroptosis and its regulator genes (FRGs) were identified to be correlated with drug resistance (Lu et al., 2017). For example, it was previously reported that liver cancer cells repressed ferroptosis by regulating the expression of *NRF2* or *MT-1G*, which promoted sorafenib resistance *in vitro* and in tumor xenograft models (Sun et al., 2016a; Sun et al., 2016b). Recently, Wang *et al*. reported that CD8+ T cells could promote tumor ferroptosis during cancer immunotherapy treatment (Wang et al., 2019). Thus, ferroptosis might play important roles during cancer progression and treatment, and a systematic study for ferroptosis and its aberrances across cancers will be helpful.

In the present study, for the first time, we performed a comprehensive analysis of genomic variations and expression profiles of the FRGs across 20 cancer types. Furthermore, we computationally modelled the ferroptosis level based on FRGs expression, and dissected the relations between ferroptosis and cancer clinical features. It was found that ferroptosis was associated with various cancer hallmarks, immune microenvironment, drug resistance and patient survival. These results highlighted the critical roles of ferroptosis in cancer and should be helpful for further investigations of ferroptosis-related molecular mechanisms and therapy development.

## Results

### Genetic alterations of ferroptosis regulator genes (FRGs) in cancers

In this study, we defined the 24 genes including *CDKN1A, HSPA5, TTC35, SLC7A11, NFE2L2, MT1G, HSPB1, GPX4, FANCD2, CISD1, FDFT1, SLC1A5, SAT1, TFRC, RPL8, NCOA4, LPCAT3, GLS2, DPP4, CS, CARS, ATP5G3, ALOX15*, and *ACSL4*, as ferroptosis regulator genes (FRGs), which were identified to play critical roles in regulating ferroptosis by previous studies (Stockwell et al., 2017). To determine the patterns for dysregulation of FRGs in cancer, we examined the genomic data, including genetic variation, somatic copy number alternation (SCNA), mRNA expression and DNA methylation data of tumor and normal tissues from 20 cancer types. The analysis of nonsynonymous mutations in 20 cancer types showed that the mutation frequencies for FRGs were generally low (Figure S1A), while 9 FRGs including *ACSL4, CARS, DPP4, GLS2, NCOA4, TFRC, FANCD2, NFE2L2, SLC7A11* in UCEC had mutation frequencies greater than 5%. Furthermore, *NFE2L2*, which encodes NRF2 and plays a master regulator role in anti-oxidant responses and has been shown to cause resistance to ferroptosis (Sun et al., 2016b), showed relatively high mutation frequencies in multiple cancers including BLCA, CESC, ESCA, HNSC, LUSC, and UCEC.

To further dissect the genetic aberrance of FRGs in cancer, the percentage of SCNA was examined and the results showed that in general SCNA occurred at high frequencies (with over five percent of all samples) in most cancer types (Figure 1A), but all FRGs in THCA showed a low frequency of SCNA. It was observed that FRGs presented diverse SCNA profiles. For example, *TTC35, HSPB1, TFRC*, and *RPL8* were more prone to copy number gain than copy number loss in almost all tumors, but *SCL7A11* and *ALOX15* showed the opposite profile. Furthermore, we analyzed the co-occurrence of mutations and SCNA between FRGs and cancer-specific oncogenes/tumor suppressor genes (TSGs) previously identified by Bailey *et al.* (Bailey et al., 2018), in which *NFE2L2* was oncogene in cancer types including BLCA, CESC, HNSC, LIHC, LUSC, and UCEC, and *CDKN1A* was as tumor suppressor gene in BLCA and LIHC. The significance was calculated via the mutual exclusivity test by DISCOVER (Canisius et al., 2016), with a false discovery rate of 1%. A total of 28 oncogenes (Figure S1B) and 43 TSGs (Figure S1C) were found to be altered with FRGs in the mutually exclusive manner in certain cancer types, and *FDFT1, RPL8, TFRC*, and *NFE2L2* were frequently exclusive to oncogenes or TSGs (Figure S1B-C).

**Figure 1.**
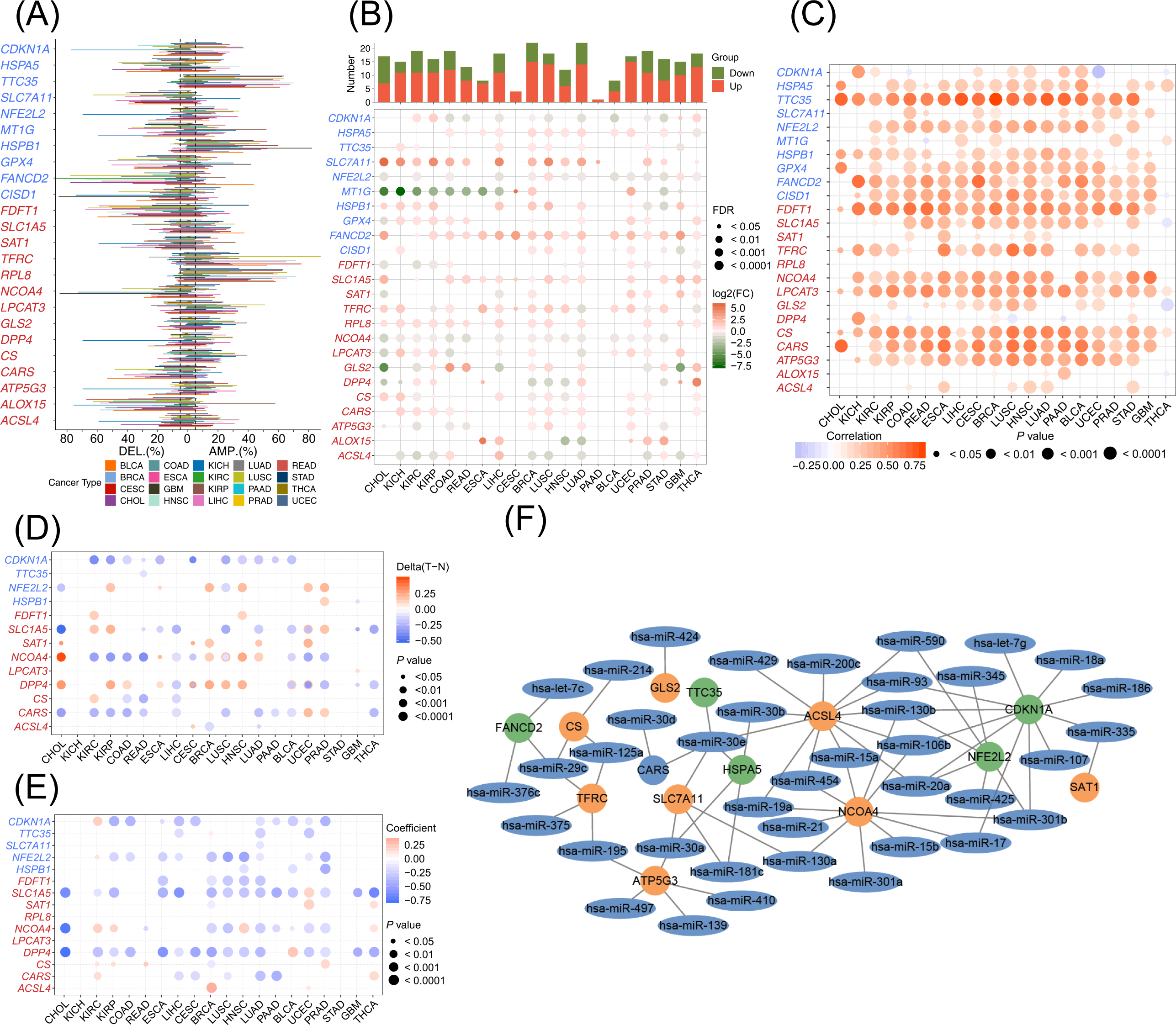
The dysregulation of the ferroptosis regulator genes (FRGs), for which the positive and negative regulators were marked in red and blue, respectively. (A) Histogram shows the frequency of somatic copy number alternations for each FRG in each cancer type. (B) Histogram (upper panel) shows the number of significant differential expressed genes, heatmap shows the fold change and FDR of FRGs in each cancer. Significantly upregulated and downregulated genes are marked in red and green, respectively. (C) The correlation between and somatic copy number alternations and expression of FRGs. (D) Heatmap shows the differential methylation of FRGs in cancers, hypermethylated and hypomethylated genes are marked in red and blue, respectively. (E) Pearson correlation of FRGs between the transcriptional expression and promoter methylation. Red and blue represented positive and negative correlation, respectively. (F) The miRNA-mRNA network for FRGs, the orange and green circle were positive and negative FRGs, respectively.

### Aberrant expression of FRGs among cancer

Besides genetic alternations, differential expression analysis was performed between tumor and adjacent normal tissues for every cancer type to investigate alteration of gene expression patterns of FRGs, and the numbers of tumor and normal sample were showed in Table 1. We found that all FRGs were differentially expressed in at least one cancer type. Several FRGs showed consistent expression patterns in cross-cancer analysis. *SLC7A11, FANCD2, CARS, SLC1A5* and *RPL8* were significantly upregulated in 15, 18, 12, 15, 14 types of cancers, respectively, while *NCOA4* was downregulated in 15 cancers (Figure 1B). Additionally, several FRGs showed miscellaneous cancer type specificities which has not been well characterized previously. For example, *HSPA5* was upregulated in most cancer types including breast invasive carcinoma (BRCA) (fold change [FC] = 1.87, adjust *P*-value = 1.42 ×10^−42^) and LUAD (FC = 1.90, adjust *P*-value = 2.45 ×10^−29^) but was significantly downregulated in THCA (FC = 0.68, adjust *P*-value = 2.67 ×10^−7^). *DPP4*, which appears to play a suppressor role in the development of cancer (Masur et al., 2006; Pro and Dang, 2004; Wesley et al., 2005), showed significant upregulation in KIRC, KIRP, ESCA, LUAD, GBM, and THCA, but was downregulated in CHOL, KICH, BRCA, LUSC, HNSC, and STAD. We also noticed that *DPP4* had opposed expression profiles in different subtypes of tumors in the lung and kidney. This demonstrated that FRGs might play different roles in different cancers.

**Table 1.**
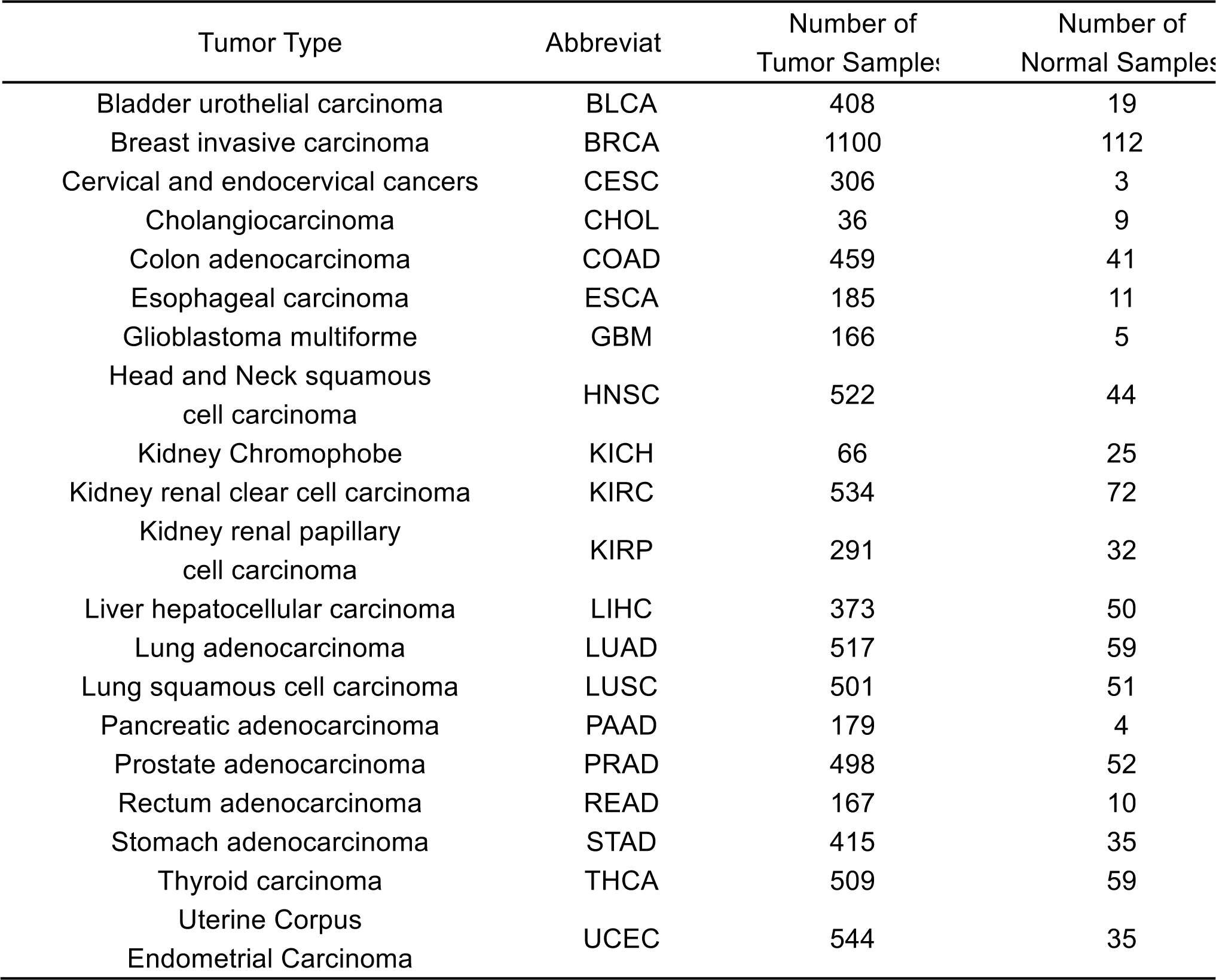
The abbreviations and numbers of samples for the 20 types of tumors investigated in this study.

Since SCNA in tumors plays a critical role in regulating gene expression, we evaluate the effects of SCNA on gene expression of FRGs. The Pearson correlation between gene expression and the copy number from masked copy number segment of TCGA was examined. The results showed that the expression of most FRGs was obviously correlated to SCNA in most tumors (Figure 1C). For example, the expression of citrate synthase (*CS*), which participates in oxidative metabolism, was significantly associated with SCNA in all cancer. This result indicates that the aberrance of copy number for FRGs is common in most cancers and can have an influence on gene expression.

Besides SCNA, the methylation of promoter can regulate gene expression, and aberrant DNA methylation of the promoter is associated with tumorigenesis (Shen and Laird, 2013). We observed that FRGs showed complex methylation patterns in the 20 cancer types (Figure 1D), only *CDKN1A* showed hypomethylation in 11 tumors consistently. For example, we observed that *DPP4* showed hypermethylation in seven cancer types and hypomethylation in four cancer types. Although there were differences in the pattern of methylation for FRGs, a negative relationship was observed between gene expression and DNA methylation overall (Figure 1E). This result demonstrated that promoter DNA methylation may regulate the expression of FRGs in tumors.

Besides SCNA and DNA methylation, microRNAs (miRNAs) can regulate gene expression and become involved in cancer development (Macfarlane and Murphy, 2010). To determine which miRNAs are involved in regulating ferroptosis, we performed the analysis to reveal the network of miRNA-FRGs. The starBase database was used to infer miRNA that potentially targeting FRGs, and those miRNAs that were significantly negatively correlated with gene expression were thought to be involved in the regulation of FRGs (Li et al., 2014). As the network showed (Figure 1F), all of the interactions between miRNA and FRGs occurred in more than six cancers. It is obvious that FRGs could be targeted by miRNA with high frequency, including *ACSL4, NCOA4*, and *CDKN1A* (Figure 1F). To further understand the aberrances of the miRNA-FRG network in tumors, we conducted differential expression analysis of miRNA and counted the expression aberrances of miRNA across cancer types (Figure S1D). It was found that several miRNAs had consistent expression trends in tumors, for example, has-miR-93 targeted *CDKN1A* and was downregulated in 11 cancers. On the other hand, miRNA had different expression trends in different tumors. For example, has-miR-375 targeted *ACSL4* in five tumors, but it showed upregulation in two cancers and downregulation in three cancers. This suggests that miRNA play a regulatory role in FRGs expression, and the aberrant expression of miRNA in the tumors could impact ferroptosis.

Since SCNA, DNA methylation, and miRNA all can regulate FRGs expression, but the contribution of a single factor on gene expression is not clear. The linear regression approach was applied to analyze the contribution of each factor and diminish the confounding effects. As the results showed (Figure S1E), SCNA was positively correlated with gene expression, while methylation and miRNA showed a negative correlation. We further observed multiple regulated patterns in FRGs, and the expression of several FRGs was only related to a single factor in several tumors, while others were related to multiple factors. For example, *ACSL4* expression was only associated with miRNA in seven cancers, including BLCA, CESC, KIRC, KIRP, LUAD, PRAD, and UCEC. SCNA played the only significant regulator role for *CARS* in CHOL, ESCA and colorectal cancer. However, the expression of *NCOA4* and *NFE2L2* was significantly regulated by three factors in seven and ten tumors, respectively. Thus, all the FRGs show diverse regulation patterns in different cancers. This suggests that the expression regulation patterns of all FRGs were tumor-specific.

### Computational modeling the ferroptosis level among cancers

To further understand the role of ferroptosis in tumorigenesis and investigate the factors or biological processes associated with ferroptosis, the ferroptosis potential index (FPI) was modeled based on the enrichment score (ES) of positive core machine components calculated by ssGSEA minus that of negative core machine components. We evaluated the FPI through three independent GEO gene expression datasets of tumor cell lines that were treated with erastin or withaferin A (WA), which were reported as an inducer of ferroptosis (Dixon et al., 2012; Hassannia et al., 2018), or ferrostatin that was identified as an inhibitor of ferroptosis (Zhang et al., 2019). The FPI was calculated for the gene expression datasets in neuroblastoma cells (GSE112384), clear cell carcinoma cells (GSE121689) and liver cancer cells (GSE104462) (Figure S2). The results showed that erastin and WA increased the FPI remarkably in all of this three cell lines, while ferrostatin obviously decreased the FPI compared to control group (Figure S2A, S2C, S2E). Because the increases of the *CHAC1* and *PTGS2* mRNA and ACSL4 protein were associated with cells undergoing ferroptosis, but these changes were not consistent in all experiments (Stockwell et al., 2017), we compared the mRNA expression of these three genes in this three cell lines. As results shown, the mRNA expression of *PTGS2* could not significantly distinguish the ferroptosis status in these experiments. For *ACSL4*, there was no obvious changes in cell lines with ferroptosis induced by erastin or WA, although decrease was observed in the ferroptosis-inhibited cells (Figure S2B, S2D). *CHAC1* was upregulated in ferroptosis-induced cell lines treated with erastin or WA, but only slightly upregulated in ferroptosis-inhibited cells (Figure S2F). Thus, the FPI could be used to represent the potential level of ferroptosis based on the transcriptome data.

Based on the computed marker of ferroptosis, we compared the differences of FPI between tumor and normal tissues with the cancer genome atlas (TCGA) data (Figure 2A). Of note, significant differences were found in most cancers such as lung cancer and gastrointestinal cancer, and higher FPI was observed in most tumors except for BRCA and UCEC. We also noticed that the FPI of normal tissues in female cancers (BRCA, UCEC, and CESC) was higher than most tumor/normal samples in all the cancers. To further dissect the factors involved in these different FPI patterns, we examined the FPI for different subtypes of BRCA. The results presented in Figure 2B showed that all the estrogen receptor (ER)-positive, progesterone receptor (PR) positive and HER2 positive patients had lower FPI than the negative patients, respectively. Furthermore, triple negative breast cancer (TNBC) samples showed higher ferroptosis level than non-TNBC samples (Figure 2C), which was consistent with previous study (Kettner et al., 2016). Furthermore, among the different subtypes of kidney cancer, the FPI of tumor samples was higher than normal in KIRP and KIRC, while the FPI in KICH was lower (Figure 2A). In addition, to further dissect the variances of ferroptosis level in different histological types, the FPI in the esophagus squamous cell carcinoma (ESCC) and esophagus adenocarcinoma (ESAD) were compared. The results showed that FPI in ESCC was significantly higher than ESAD and normal samples, while ESAD was higher than normal but not significant (Figure 2D).

**Figure 2.**
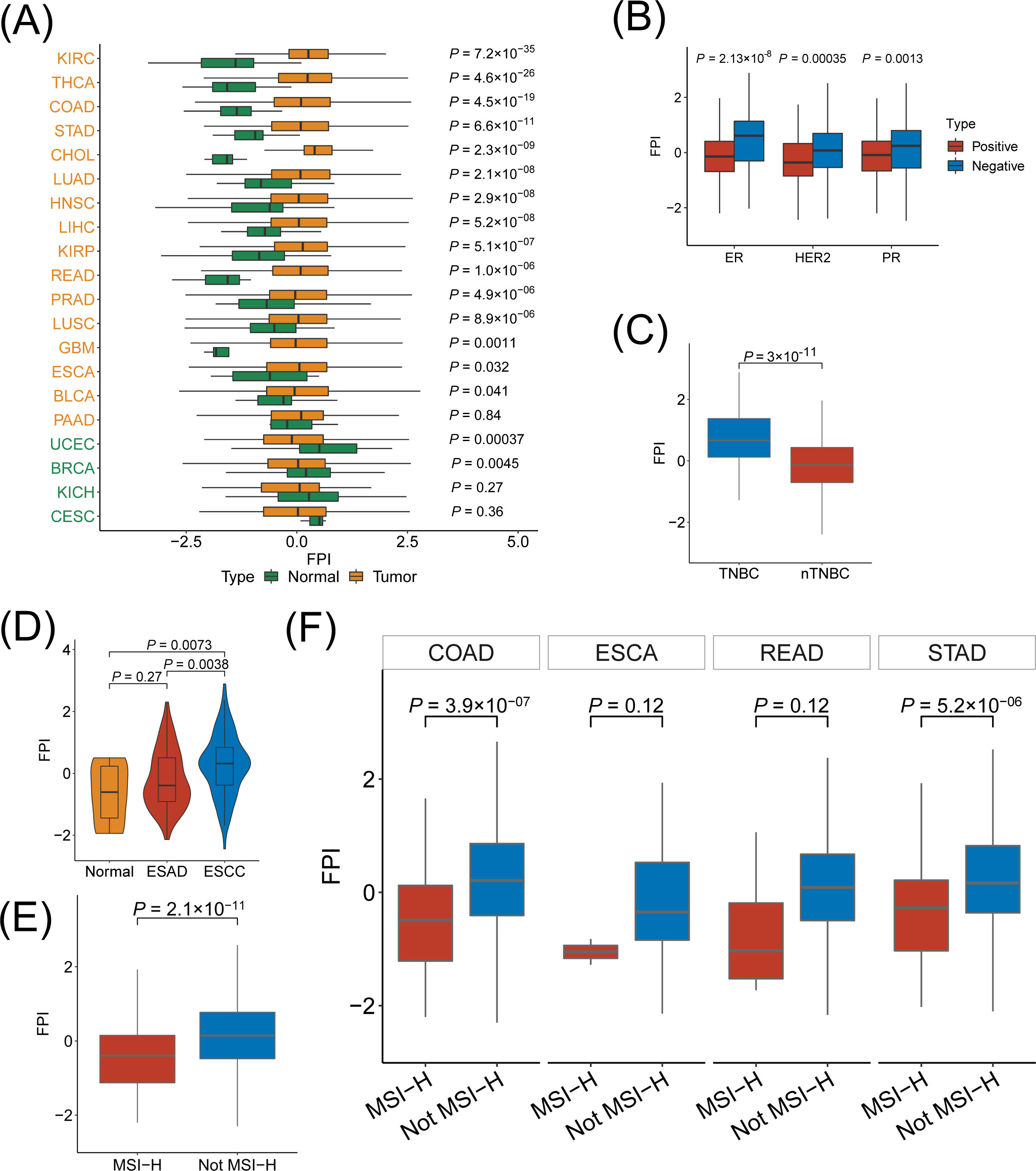
The relations between FPI and histological types and molecular subtypes among cancers. (A) The different FPI between tumor and normal tissues among cancers. (B) Box shows the difference of FPI between positive and negative receptor in breast cancer. (C) The difference of FPI between TNBC and nTNBC. (D) The different FPI among different histological types of esophageal carcinoma. The FSI for different MSI status of overall (E) and detailed (F) digestive system neoplasm.

Besides the cancer types and subtypes, we analyzed the correlation between the FPI and remarkable molecular features such as microsatellite instability (MSI) and driver gene mutations. The FPI decreased obviously in tumors with high-level of MSI (MSI-H) compared to non-MSI-H tumors in gastrointestinal tumors (Figure 2E). The FPI was remarkably lower in COAD and STAD patients with MSI-H, and slight decrease was also observed in ESCA and READ (Figure 2F). Furthermore, we analyzed the correlation between FPI and 375 driver genes that frequently mutated in tumors based on regression-based analysis. Through controlling for cancer type and tumor mutation burden (TMB), 81 genes were found to be correlated, and most of them were negatively associated with FPI (adj.*p* < 0.1, Figure S3A). Strikingly, somatic mutations, namely *NFE2L2, BRAF*, and *TP53*, were positively associated with FPI in pan-cancer associations, but a mutation in *TP53* was negatively correlated with ferroptosis in LUSC. This observation was verified by the comparison of FPI between the *TP53* mutant and wild type groups (Figure S3B), consistent with previous finding that showed that *TP53* played a dual role in regulating ferroptosis (Kang et al., 2018). The influences of *KRAS* mutations on FPI were also investigated, and the results showed that the relation between *KRAS* mutation and FPI was significantly negative in gastric tumors (Figure S3C) but slightly positive in hepatocellular carcinoma (Figure S3A, S3C). These results indicated that the tumor type and molecular context were critical for the regulation of ferroptosis, which was consistent with previous study (Tsoi et al., 2018). Furthermore, the differential expression of FRGs between wild type and mutant tumors for *TP53* (Figure S3D) and *KRAS* (Figure S3E) were found to be ubiquitous in most cancers. Taken together, ferroptosis was negatively related with MSI-H in cancer.

### Association between ferroptosis and pathways in cancer

To further elucidate the association between the FPI and other genes and pathways, we calculated Spearman correlation coefficients between FPI and all the genes including FRGs (Supplementary Table1). It was found that generally FPI was positively and negatively correlated with the expression of positive and negative FRGs, respectively, while various exceptions were also observed (Supplementary Table1). The related cellular signaling of ferroptosis in cancer was investigated by gene set enrichment analysis (GSEA) for each cancer based on the transcriptome of two tumor groups with the top and bottom thirty percent of FPI. It was observed that metabolism related pathways in KEGG usually enriched in the tumor with lower FPI, and pathways frequently enriched (>6 cancers) were presented in Figure 3A. For example, terpenoid backbone biosynthesis and steroid biosynthesis were enriched in the low FPI group in 19 and 16 cancers, respectively (Figure 3A). Peroxisome, biosynthesis of unsaturated fatty acids, etc. were also significantly correlated with lower FPI in many cancer types with no opposite was observed (Figure 3A). Furthermore, the relations between cancer hallmarks and FPI were also analyzed, and the results showed that 15 hallmarks were frequently significantly correlated with FPI (Figure 3B). For example, KRAS signaling, epithelial-mesenchymal transition, IL6 JAK STAT3 signaling, WNT Beta-catenin signaling were enriched in the high FPI group, which indicated that ferroptosis was positively related with these oncogenic pathways (Figure 3B). Also, metabolism related hallmarks were observed to be negatively related with ferroptosis, which was consistent with the pathway analysis (Figure 3B).

**Figure 3.**
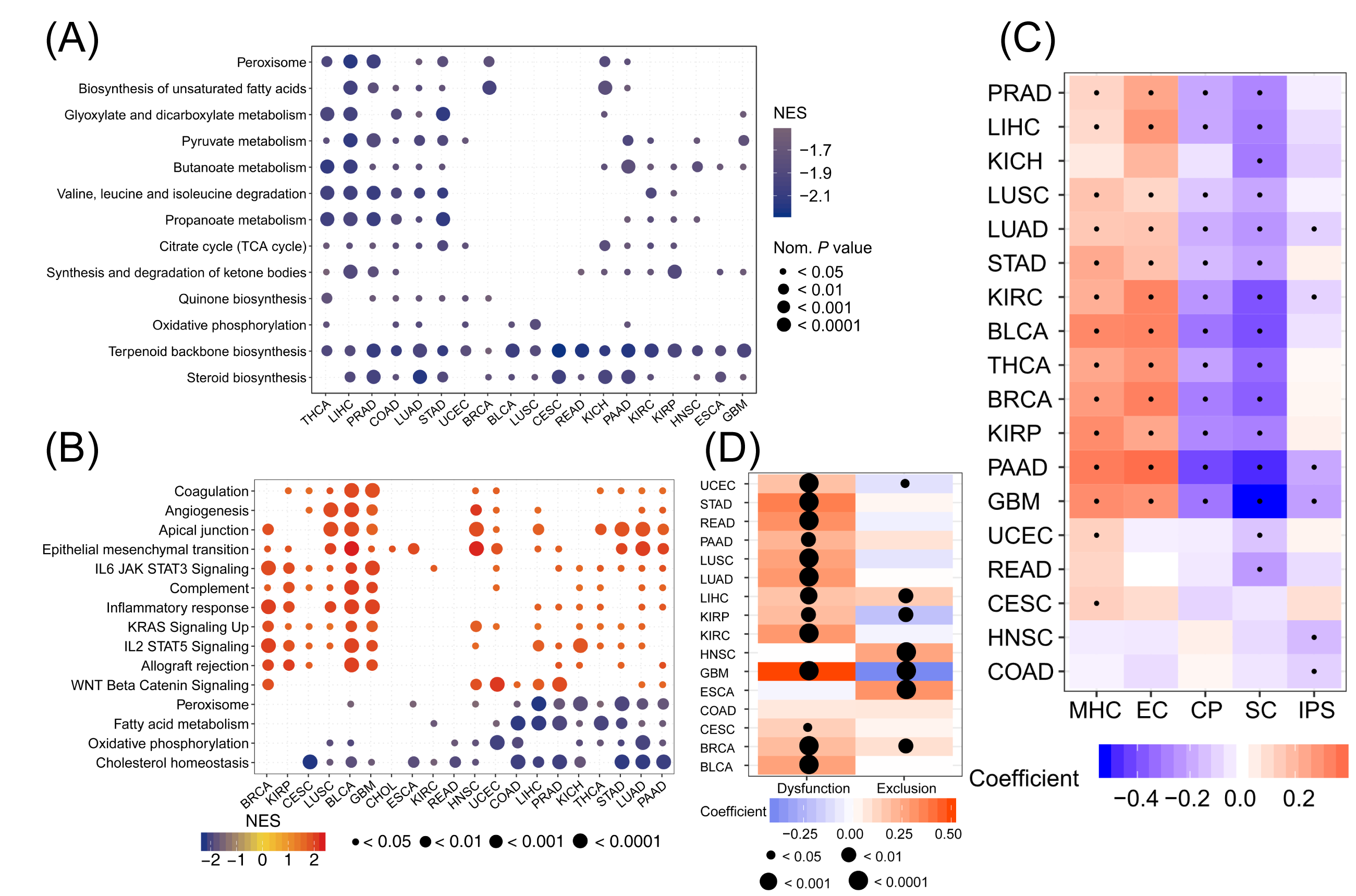
Relations between the ferroptosis and signaling pathways and immunophenotypes. Enrichment analysis for metabolism pathway (A) and cancer signaling (B) between high- and low-FPI tumor tissues. NES is the normalized enrichment score in the GSEA algorithm. (C) Correlations between immunophenotypes and the FPI, and dots indicate statistically significant results (*P* value<0.05).

Since previous studies showed that ferroptosis was related to the immune response process (Matsushita et al., 2015), here we investigated the association between ferroptosis and immune microenvironment in tumor. The results showed that in several cancers FPI was weakly negatively correlated with immunophenoscore (IPS) (Figure 3C), which could predict the response of the immune checkpoint blockade in melanoma tissue (Charoentong et al., 2017). To better understand the relationship between ferroptosis and the response of immunotherapy, we calculated the Spearman coefficients and found the FPI was positively correlated with dysfunction score of T cells of 13 cancers. But no consistent effect of the FPI on the exclusion score of T cells was observed (Figure 3D). Furthermore, FPI was positively correlated with MHC and effector cells (EC) but negatively correlated with immunomodulators (CP) and suppressor cells (SC) in most cancers (Figure 3C). The associations between ferroptosis and immune cell types were further evaluated in detail, and the results showed that FPI showed a positive correlation with macrophages in most cancers and a negative association with the activated dendritic cells, activated mast cells, plasma cells and T follicular helper cells (Figure S3F). Thus, there might be close connections between ferroptosis and immune microenvironment and function of T cells, and more investigations were needed to reveal the details.

### Clinical relevance of ferroptosis and its regulators

Since ferroptosis was involved in cellular metabolism and immune microenvironment, it was proposed that ferroptosis and its regulators should be correlated with cancer survival and other clinical characteristics. Based on the clinical data from TCGA, the survival analysis was performed and the results were consistent since it is found that the lower FPI predicted a better survival in five cancers including GBM, KIRC, KIRP, LIHC, and LUAD (Figure 4). We further dissected the factors that might impact FPI including tumor stage, cigarette smoking, age, and gender. As the results shown (Figure S3G), tumor stages were positively correlated with FPI in LIHC, KIRC, KIRP, and COAD. Only the FPI in LUAD tumors showed a positive correlation with cigarettes smoked per day. Age was positively associated with FPI in GBM and PRAD but showed a negative correlation in seven cancers such as BRCA and LIHC. It was observed that FPI was remarkably different between genders (Figure S3H), FPI for female tumor samples was higher than male tumor samples in KIRC, LIHC, LUAD, and LUSC, but lower in STAD. Significant difference in FPI were also observed among different races in BLCA, BRCA and ESCA. Furthermore, we found that the FPI was correlated cancer metastasis, recurrence, primary and follow outcome in several cancers. In addition, we evaluated the association between FPI and other clinical characteristics including alcohol abuse, fatty liver, hemochromatosis, and viral infection (HPV, HBC, HCV, and EBV), while no significant results were observed. These results indicate that ferroptosis might play critical roles in cancer survival, and clinical factors have tumor-specific effects on FPI.

**Figure 4.**
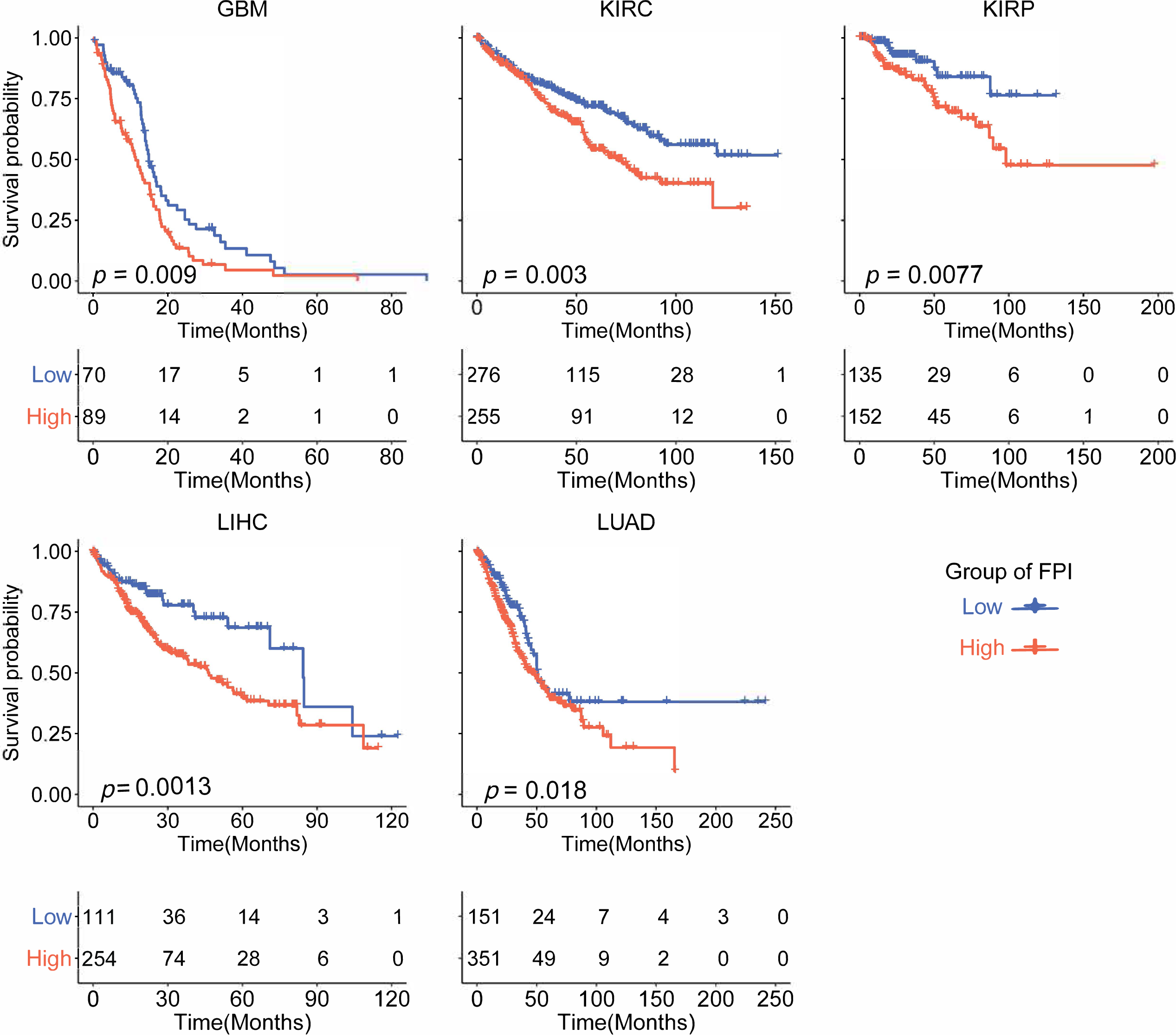
Kaplan–Meier analysis of overall survival according to the FPI among cancers.

To further dissect the clinical relevance of ferroptosis in cancer, the roles of FRGs in cancer survival were analyzed. The survival analysis showed that all FRGs were associated with overall survival in at least six cancer types (Figure S3I), but the correlations were miscellaneous since most FRGs could both serve as protective and risk factors in different cancers. For example, patients with higher expression of *FANCD2* showed better survival in kidney cancer, LIHC, and LUAD, but showed worse survival in READ, STAD, and BLCA (Figure S3I). However, high expression of *CARS* and *MT1G* showed consistent better survival among cancers including KICH, UCEC, PAAD, and HNSC (Figure S3I). Thus, the functional roles of FRGs in cancer survival should be further explored.

To study the potential clinical implications of ferroptosis, we examined the association between the gene expression of FRGs and 156 clinically actionable genes (CAGs), including 136 genes for targeted therapy and 20 genes for immunotherapy across cancer types (Mak et al., 2016; Van Allen et al., 2014). As a FRG, *CDKN1A* regulates iron uptake and *GPX4* abundance (Stockwell et al., 2017), was also a CAG targeted by anti-cancer drugs, such as the inorganic compound arsenic trioxide (Hassani et al., 2018). We found that all 136 CAGs and FRGs had significant co-expression relationships that were significantly different across cancers as shown in Figure S4A. The number of CAGs correlated with FRGs ranged from 45 in LUSC to 132 in THCA (Figure S4B). The FRG-CAG correlation pairs ranged from 69 in LUSC to 581 in THCA (Figure S4C). For example, *GPX4* co-expressed with *MAP2K2* in 20 cancer types, and showed a significantly positive correlation with *TNFRSF4*, which are targeted genes for immunotherapy. *ACSL4* is significantly correlated with 17 of 20 genes targeted for immunotherapy, suggesting that *ACSL4* has a potential effect on cancer immunotherapy (Figure S4D). Furthermore, besides co-expression, the protein-protein interactions among drug-targeted CAGs and FRGs were analyzed, and the results presented in Figure S4E showed that the interactions were obvious. For example, *EGFR*, the target of lapatinib, gefitinib and afatinib, was positively associated with FRGs including *DPP4* (R = 0.15) and *HSPA5* (R = 0.31), and negatively correlated with *FDFT1* (R = -0.24), while there were protein-protein interactions between EGFR and DPP4, HSPA5, FDFT1 (Figure S4E). These results indicated that ferroptosis might be involved in drug effects by interacting with targeted clinically actionable genes. Taken together, our studies suggest that clinically actionable genes are closely related to FRGs, which highlights the significance of ferroptosis in cancer treatment, including both immunotherapy and targeted therapy.

To further understand the correlation between ferroptosis and drug sensitivity, the area under the percent viability curve (AUC) approach was employed to evaluate drug sensitivity (Basu et al., 2013) and calculate the correlation between FPI and drug sensitivity across cancer cell lines. Drugs associated with FPI were also tested for their correlation with FRG expression. We identified that FPI was significantly associated with 64 drugs, including 12 that were negatively correlated and 52 that were positively correlated (Figure S4F). To explore the effects of each FRG on drug sensitivity, we also analyzed the associations between the drug sensitivity of 64 cancer drugs and the expression of FRGs, and found 521 significantly correlated pairs (Figure S4F). Among them, the expression of *FANCD2* was correlated with AUC in 57 drugs, while no significant relation was observed for *CISD1* (Figure S4F). Furthermore, the associations classified the FRGs into two group, one group included genes such as *FANCD2* and *ATP5G3*, which were positively associated AUC of docetaxel, trametinib, etc. (Figure S4F). The other group included *SLC7A11, SAT1, HSPA5*, etc., which were negatively associated (Figure S4F). Taken together, these results suggest that ferroptosis was correlated to the sensitivity of multiple drugs.

## Discussion

Ferroptosis is driven by damage of cell membranes caused by the loss of activity of GPX4 (Feng and Stockwell, 2018). Researchers identified that ferroptosis is closely related to tumorigenesis and plays an important role in cancer treatment (Shen et al., 2018; Yang et al., 2014). However, there is a lack of systematic studies on ferroptosis and its regulator genes across cancer types. In this study, we employed multi-omics data and clinical data across 20 cancer types from TCGA and revealed the global alterations of ferroptosis regulator genes at genetic, epigenetic and transcriptional levels. We also used ssGSEA to process expression data to establish FPI to characterize ferroptosis and addressed which genetic and nongenetic factors (including drug and patient phenotype) were related with FPI. Different molecular types affected the ferroptosis in breast cancer and gastrointestinal cancer, which meant that the responses of different molecular subtypes to treatment may be related to ferroptosis. On the other hand, molecular subtypes need to be considered when ferroptosis is applied as a therapeutic strategy.

The mechanism by which ferroptosis regulates tumor cell growth and proliferation is still unclear, but the relationship we observed between FPI and hallmarks of cancer could improve the understanding the role of ferroptosis. The results of GSEA demonstrate that the level of ferroptosis is closely related to tumor-associated hallmarks in most cancers. Ferroptosis genes can play both oncogene and tumor suppressor roles in cancer (Kang et al., 2018), and the FPI acts as a protective or risk factor across cancer types. We also found that several common clinical factors affected ferroptosis, such as cigarette smoking, BMI and tumor stage. As we know, cigarette smoking is a risk factor for esophageal cancer, lung cancer and kidney cancer (Chow et al., 2010; Mao et al., 2011; Pesch et al., 2012). In our results, the number of cigarettes per day was positively correlated with the ferroptosis potential index in LUAD, which may be because the cigarette-induced oxidation reaction promoted lipid peroxidation (Barreiro et al., 2010; Guan et al., 2013; Louhelainen et al., 2008). FPI also varied between the sexes in several cancers, including LIHC, LUAD, LUSC, and STAD and between different races in BRCA, BLCA, and ESCA, which implied that gender and race need to be considered when using ferroptosis as a treatment strategy. We also noticed that better clinical outcomes or status in several cancer types also have lower FPI, which further confirmed the dual role of ferroptosis. Thus, a different strategy of regulating the ferroptosis of tumor cells may benefit patients and improve prognosis.

Furthermore, we showed that FRGs were co-expressed with most clinically actionable genes and interacted with genes modulated by drugs, suggesting that studying ferroptosis may improve the strategy for cancer therapy. We observed that the co-expression of FRGs and CAGs could be roughly divided into two groups, and drugs that target clinical genes could have complex regulatory effects on ferroptosis. Most strikingly, the expression of *GPX4* was associated with *MAP2K2* in 20 cancers, which indicated that clinically actionable gene *MAP2K2* may play an important role in ferroptosis. To further explore drug sensitivity and ferroptosis, we observed that the FPI could characterize the sensitivity of many drugs, the AUC of many drugs was inversely associated with the FPI in cancer cell lines, which implied that regulating the ferroptosis of tumor cells may improve the therapeutic effect of cancer treatments. However, opposite effects were observed regarding the effects of drugs on ferroptosis. Thus, further detailed studies should be carried out to determine the functions and mechanisms of ferroptosis in different cancers. In the present study, we presented a systematical analysis of ferroptosis and its regulator genes across cancers. Most FRGs were aberrantly expressed in tumors among various cancer types, while frequent CNA and differential DNA methylation contributed greatly. We established the FPI to evaluate the ferroptosis level and found that the FPI were higher in tumors than in the adjacent normal tissues in most cancers and were associated with clinical features, and cancer metastasis, recurrence, outcome and drug sensitivity. These findings highlighted the potential of ferroptosis-based cancer therapy.

## Limitations of the Study

Since current omics data only provide RNA-level quantifications for FRGs while ferroptosis process rely on proteins, although we tried to infer the ferroptosis status precisely, there should be a variety of inaccuracy. Furthermore, the detailed molecular mechanisms and genes involved in ferroptosis are still to be dissected, our study based on current knowledge might need further update and improvement due to new discoveries.

## Methods

### Datasets and Source

The mRNA expression data, copy number alteration thresholded data, masked copy number segmentation data, and DNA methylation 450K data of twenty cancers, including bladder urothelial carcinoma (BLCA), breast invasive carcinoma (BRCA), cholangiocarcinoma (CHOL), colon adenocarcinoma (COAD), cervical and endocervical cancers (CESC), esophageal carcinoma (ESCA), glioblastoma multiforme (GBM), head and neck squamous cell carcinoma (HNSC), kidney chromophobe (KICH), kidney renal clear cell carcinoma (KIRC), kidney renal papillary cell carcinoma (KIRP), liver hepatocellular carcinoma (LIHC), lung adenocarcinoma (LUAD), lung squamous cell carcinoma (LUSC), pancreatic adenocarcinoma (PAAD), prostate adenocarcinoma (PRAD), rectum adenocarcinoma (READ), stomach adenocarcinoma (STAD), thyroid carcinoma (THCA), uterine corpus endometrial carcinoma (UCEC), which had both tumors and normal samples were downloaded from Firehose (http://gdac.broadinstitute.org). Mutation data, miRNA-seq data, and clinical data were downloaded from the Xena Browser (https://xenabrowser.net/datapages/). Additional gene-centric RMA-normalized gene expression profiles and drug response data of over 1000 cancer cell lines were accessed from the Genomics of Drug Sensitivity in Cancer (GDSC) database (https://www.cancerrxgene.org/downloads) (Yang et al., 2013). Immune associated data, including immune cell type fractions and immunophenoscore were obtained from TCIA (https://tcia.at/home) (Charoentong et al., 2017). Integrated protein-protein interaction data was obtained from the Human Protein Reference Database (http://www.hprd.org/) and BioGRID (https://thebiogrid.org) (Keshava Prasad et al., 2009; Peri et al., 2003). We obtained clinically actionable genes (CAGs) from a previous study (Van Allen et al., 2014) (https://software.broadinstitute.org/cancer/cga/target).

### Differential expression analysis of mRNA

To test genes differentially expressed between tumor and normal tissue, gene expression data for 20502 genes across 20 cancer types were downloaded from TCGA at FireBrowse (http://gdac.broadinstitute.org, 2016 January). Then, the fold change and adjusted *P*-value were calculated by the edgeR package (Robinson et al., 2010). We defined genes with an adjusted *P*-value less than 0.05 as the differential expression genes (DEGs).

### Establishing the Ferroptosis Potential Index Model

The index to represent the ferroptosis level was establish based on the expression data for genes of ferroptosis core machine including positive components of *LPCAT3, ACSL4, NCOA4, ALOX15, GPX4, SLC3A2, SLC7A11, NFE2L2, NOX1, NOX3, NOX4, NOX5* and negative components of *FDFT1, HMGCR, COQ10A, COQ10B*. The enrichment score (ES) of gene set that positively or negatively regulated ferroptosis was calculated using single sample gene set enrichment analysis (ssGSEA) in the R package ‘GSVA’ (Hanzelmann et al., 2013), and the normalized differences between the ES of the positive components minus negative components was defined as the ferroptosis potential index (FPI) to computationally dissect the ferroptosis levels/trends of the tissue samples.

### Somatic Copy-number Alteration (SCNA) and Mutation Analysis

The heterozygosity and homozygosity of amplification and deletion were included to elevate the copy-number alteration of each gene, in which over five percent was regarded as high-frequency SCNA. Pearson’s correlation between expression values and copy number segment values of each gene was calculated to evaluate the association between CNA and expression. The R package “DISCOVER” was employed to evaluate mutual exclusivity between FRGs and tumor suppressor genes or oncogenes across tumor samples in each cancer type (Canisius et al., 2016). The mutation and CNA events were integrated, while only homozygous amplification and deletion were included, and only protein-coding mutations were retained. For each cancer type, the genes were considered to be mutually exclusive if they had a q value of 0.05.

### DNA methylation analysis

The R package “IlluminaHumanMethylation-450kanno.ilmn12.hg19” from Bioconductor was imported to annotate the methylation probe for the promoter of each gene. Differential methylation of each gene in tumor and normal samples was tested by the Wilcoxon signed rank test, and genes that were significantly hypomethylated or hypermethylated were identified using a *P*-value cutoff of 0.05. Pearson’s correlation between the transcriptional expression of FRGs and the Beta value of the promoter DNA methylation were calculated and considered significant if the *P*-value < 0.05.

### miRNA expression analysis

To investigate the mechanisms of dysregulation for ferroptosis regulator genes in cancer, we searched potential miRNAs which might regulate the FRGs based on miRNA-target intersections in starBase. The Spearman correlation between the expression of miRNA and FRGs was statistically evaluated (adjusted *P*-value < 0.1, rho < -0.1). Cytoscape software was used to visualize the high-frequency interaction networks among FRGs and miRNA (Shannon et al., 2003).

### Multivariate Regression Analysis of Gene Expression

To assess which factors had significant effects on FRG expression, the expression of each FRG was modeled by linear regression as a function of the median miRNA expression, the median Beta value of promoter methylation, and the copy number of the genes.

### Clinical Features Analysis

The R package “survival” was used to assess the prognosis potential of the FRGs and ferroptosis potential index among cancers. For survival analysis, the expression threshold was exhaustively tested and the one with most significant *P*-value was considered the best cut-off. To test the association between ferroptosis level and clinical features, the Pearson correlation was calculated between FPI and tumor stages, age, body mass index (BMI) and cigarette exposure per day, which were converted to numeric variables (“stage1” = 1, “stage2” = 2, etc.). Wilcoxon rank sum tests and Tukey’s tests were used to determine the impact on the ferroptosis potential index for other clinical characteristics including race, remission status, and alcohol. The influence of different MSI statuses(Liu et al., 2018), histologic types, and molecular subtypes on FPI were also considered(Ciriello et al., 2015).

### Immune Features Analysis

To study the relationship of ferroptosis and immune microenvironments, we computed the Pearson correlation between FPI and immune parameters including immune cell fractions and 5 types of immunophenoscores.

### Identifying the FPI associated significant driver gene mutation

A total of 375 driver genes identified in previous pan-cancer research were included for analysis(Lawrence et al., 2013). To test whether a driver gene’s mutational status was significantly associated with ferroptosis among cancers, the rank-transformed FPI was modeled by linear regression as a function of the driver gene’s mutational status, ignoring the synonymous variant. To diminish the confounding effects, the rank-transformed count of the total nonsynonymous mutations and the tumor type, which were encoded as virtual variables, were included. To further characterize each cancer, the Benjamini-Hochberg method was used to correct the *P* values across 375 genes. Genes with an adjusted *P* value less than 0.05 for mutation status variable were significantly associated with FPI.

### Gene Set Enrichment Analysis

To identify the pathways associated with ferroptosis, the samples of each tumor type were divided into two groups according to the FPI, consisting of the top 30% and bottom 30%. Then, the gene set enrichment analysis (GSEA) was performed (Subramanian et al., 2005).

### Correlation between Drug Sensitivity and FPI/FRG Expression

To test the correlation between small molecular drugs and FPI and FRGs, the Pearson correlation coefficients for FPI, the expression value of FRGs and the area under the dose-response curve (AUCs) values were calculated, the results with |R| > 0.1 and *P*-value < 0.05 were considered as significantly correlated. To further investigate the influence of drugs on ferroptosis, the significant association between the expression of clinically actionable genes and FRGs were finished across cancer cell lines, and the associations were filtered with PPIs. Then the drugs which target CAGs were selected according to DrugBank (Wishart et al., 2018) (https://www.drugbank.ca).

## Acknowledgements

This work was supported by grants from National Key R&D Program of China (2018YFC1313300 to RH.X.), Program for Guangdong Introducing Innovative and Entrepreneurial Teams (2017ZT07S096 to ZX.L.), Tip-Top Scientific and Technical Innovative Youth Talents of Guangdong Special Support Program (2019TQ05Y351 to ZX.L.), CAMS Innovation Fund for Medical Sciences (2019-I2M-5-036 to RH.X.), Natural Science Foundation of Guangdong Province (2017A030313485 to RH.X., 2019A1515010634 to ZX.L.); and Science and Technology Program of Guangdong (2019B020227002 to RH.X.).

## Author contributions

ZX.L. and RH.X. designed and supervised the experiments. Z.L., Q.Z., ZX.Z., and SQ.Y. performed the data analysis with contributions from K.Y., QF.Z., X.Z., H.S., HQ.J, H.C., and F.W.. ZX.L. and Z.L. wrote the manuscript with contributions of all authors. All authors reviewed the manuscript.

## Supplemental files

**Figure S1** – Molecular alternations of FRGs. The mutation frequencies (A), mutual exclusivity between FRGs and oncogenes (B) /tumor suppressor genes (C) for FRGs among cancers. (D) The differential expressed FRGs-related miRNAs. The correlation between FRGs expression and somatic copy number alternation, DNA methylation and miRNAs.

**Figure S2** – Established the FPI in cell lines and examined the performance of the FPI. The boxes showed the FPI between withaferin A (WA), erastin or ferrostatin and the control (A, D, G). The boxes showed the comparison of three genes between stimulators and control in neuroblastoma cells (B), clear cell carcinoma cells (E) and liver cancer cells (H). The correlation between the expression of *CHAC1* and the FPI in neuroblastoma cells (C), clear cell carcinoma cells (F) and liver cancer cells (I).

**Figure S3** – The correlation between the FPI and mutation status of driver genes (A). The FPI between mutant and wild type tumors for *TP53* (B) and *KRAS* (C). The differential expression of FRGs between mutant and wild type tumors for *TP53* (D) and *KRAS* (E). The correlation between ferroptosis and immune cells (F) and clinical characteristics (G-H). The overall prognosis abilities of FRGs (I).

**Figure S4** – (A) The co-expression between FRGs and 136 genes for targeted therapy. (B) The number of CAGs correlated with FRGs. (C) The number of FRG-CAG correlation pairs. (D) The co-expression between FRGs and 20 genes for immunotherapy. (E) The interactions between drug, drug target and FRGs. (F) The correlation between expression of FRGs and the area under the dose-response curve (AUCs) for drugs.

**Table S1** - The Spearman coefficient between the FPI and genes across cancers. The genenames of the FRGs were marked in red.

