## Supplementary figures and images for "Systematic analysis of aberrances of ferroptosis reveals its potential functional roles in cancer"

### Supplemental Figure 1

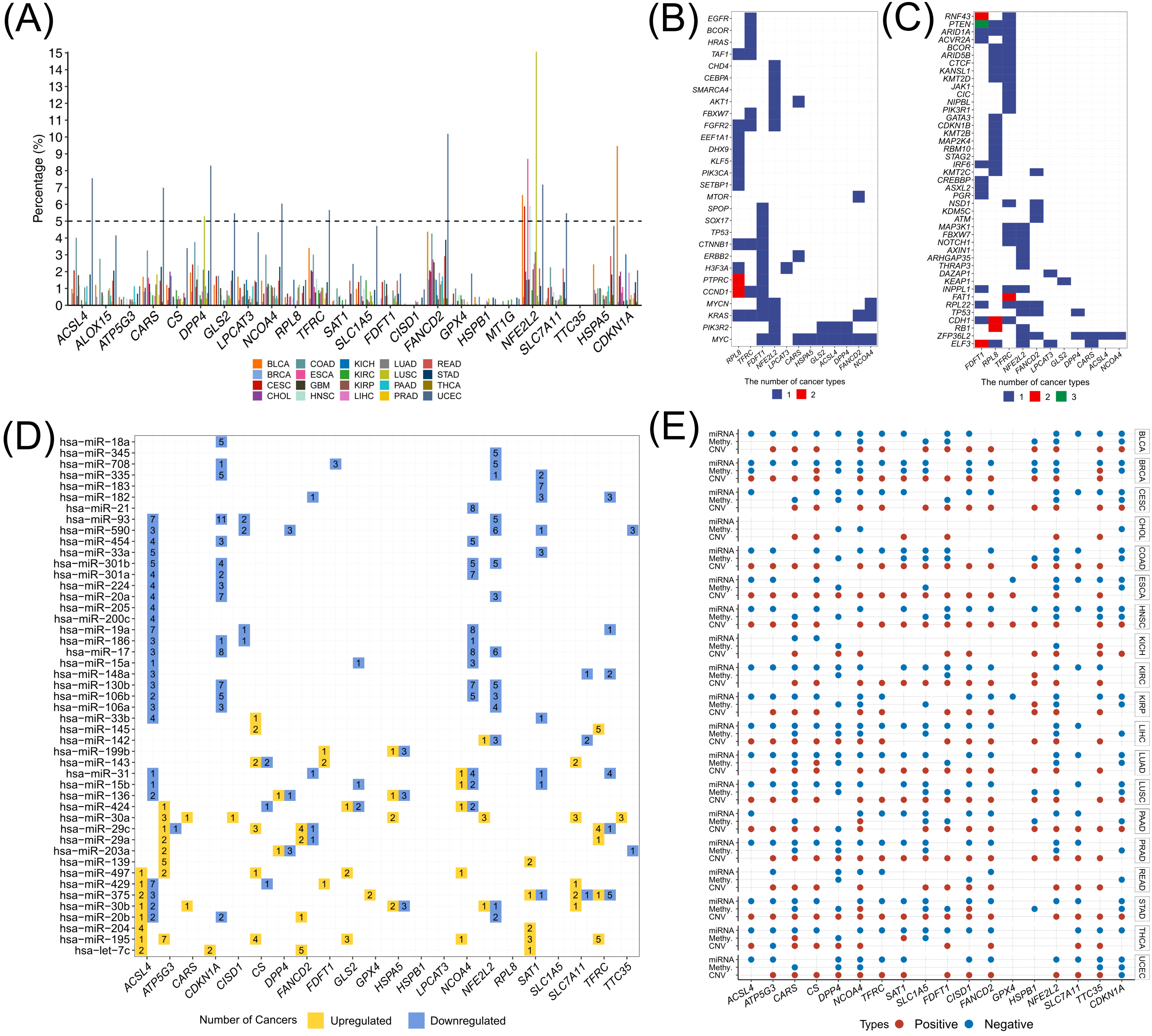

### Supplemental Figure 2

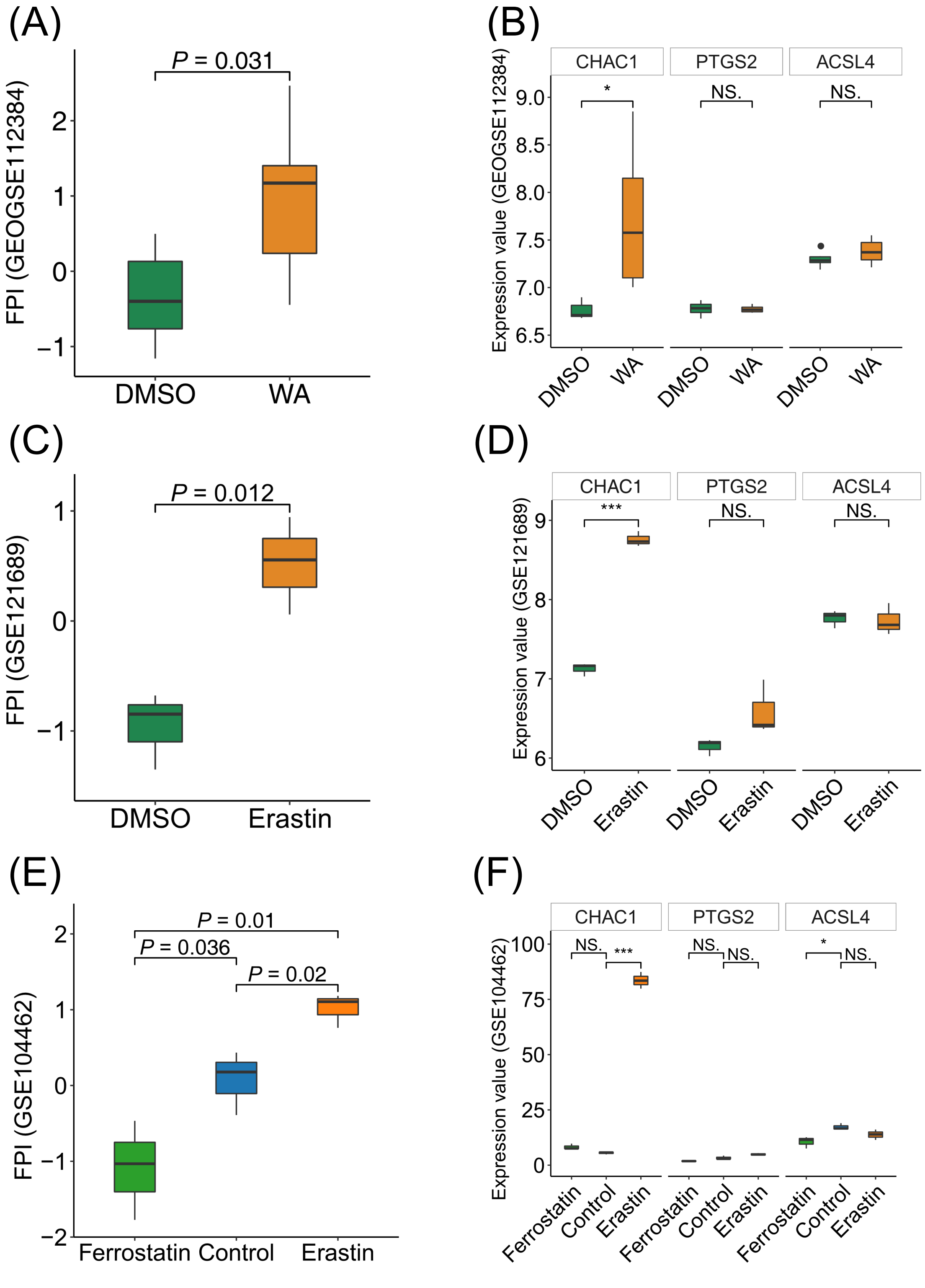

### Supplemental Figure 3

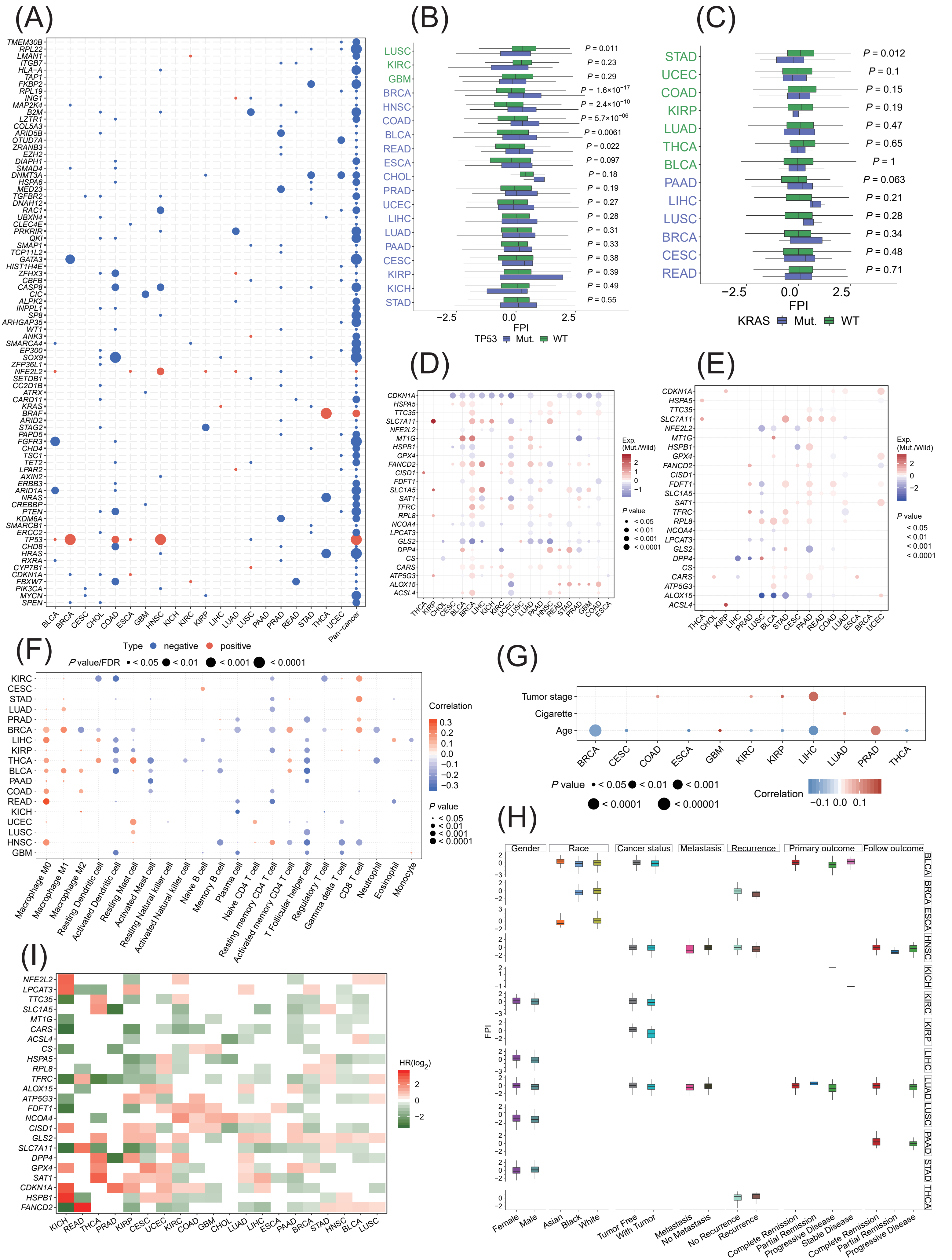

### Supplemental Figure 4

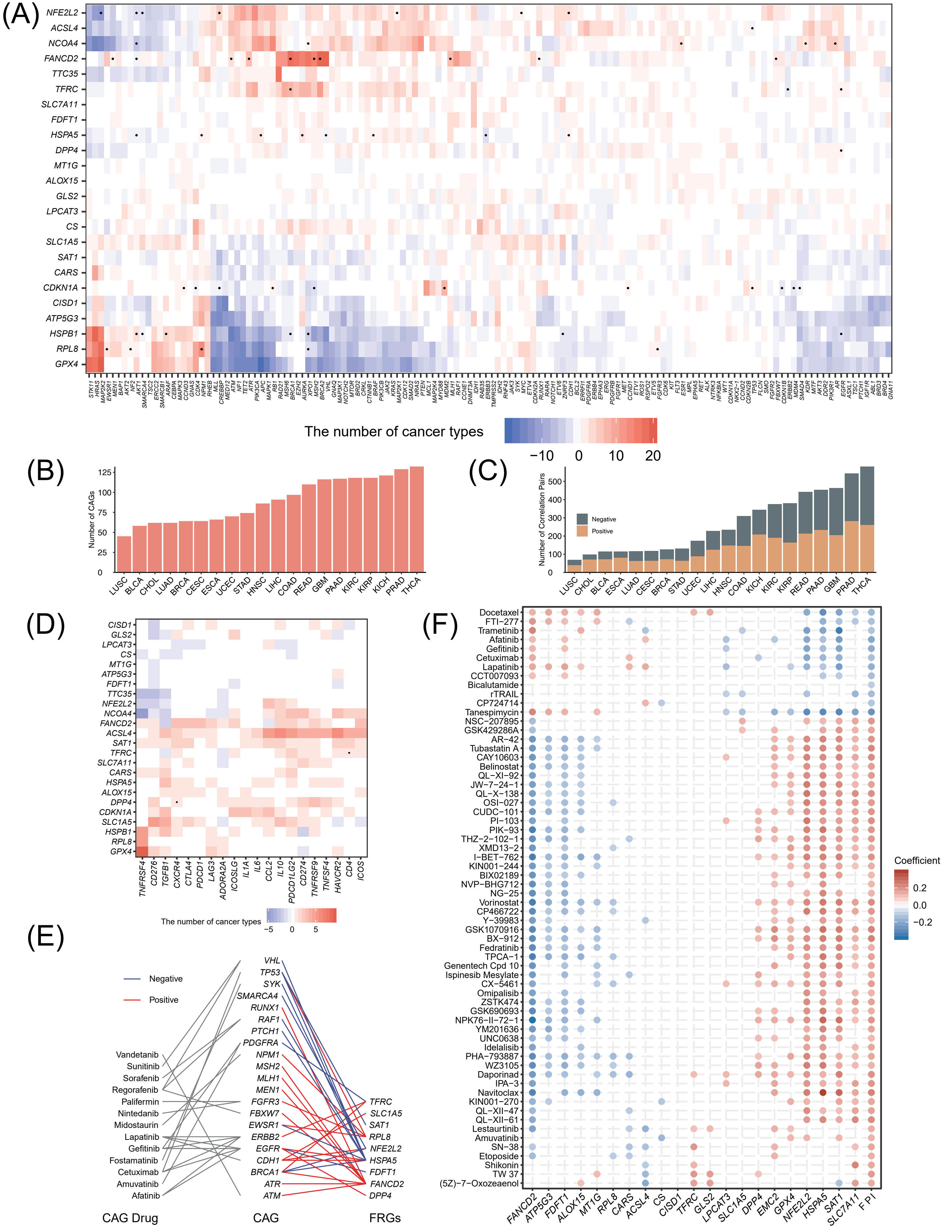
